# Robustness and universality in organelle size control

**DOI:** 10.1101/789453

**Authors:** Kiandokht Panjtan Amiri, Asa Kalish, Shankar Mukherji

**Affiliations:** Department of Physics, Washington University in St. Louis, St. Louis, MO; Department of Cell Biology & Physiology, Washington University School of Medicine, St. Louis, MO; Center for the Science and Engineering of Living Systems, McKelvey School of Engineering, Washington University in St. Louis, St. Louis, MO

## Abstract

One of the grand challenges in quantitative cell biology is understanding the precision with which cells assemble and maintain subcellular organelles. A critical property that governs organelle function is its size. Organelle sizes must be flexible enough to allow cells to grow or shrink them as environments demand, yet be maintained within homeostatic limits. Despite identification of numerous molecular factors that regulate organelle sizes we lack insight into the quantitative principles underlying organelle size control. Here we examine organelle sizes from *Saccharomyces cerevisiae* and human iPS cells with mathematical theory to show that cells can robustly control average fluctuations in organelle size. By demonstrating that organelle sizes obey a universal scaling relationship we predict theoretically, our framework suggests that organelles grow in random bursts from a limited pool of building blocks. Burst-like growth provides a general biophysical mechanism by which cells can maintain on average reliable yet plastic organelle sizes.

## INTRODUCTION

Among the most critical scales of biological organization in the eukaryotic cell is its compartmentalization into organelles. Organelle biogenesis, among the most complex tasks the eukaryotic cell performs, is the result of the coordinated synthesis of tens to hundreds of protein and lipid macromolecular species, each of which is potentially subject to stochastic fluctuations intrinsic to their production. How these molecular-scale fluctuations propagate to organelle-scale functional properties remains an outstanding question in quantitative cell biology (Chang and Marshall 2017, Vagne and Sens 2018, Sachadeva 2016, Bauer 2020). We have previously shown that organelle copy number statistics exhibit substantial cell-to-cell variability (Mukherji and O’Shea 2014), though the root mechanisms of this variability remain subject of much debate (Craven 2016, Choubey 2019). Here we turn our attention to the ability of the eukaryotic cell to control a closely linked biophysical property that determines organelle function: organelle size.

Pioneering work on a wide variety of organelles has focused on characterizing average organelle sizes and begun to unravel the molecular mechanisms underpinning this size control (Marshall 2016). The physiological importance of controlling organelle size is highlighted by their massive expansion upon increased demand for their outputs. The volume of the endoplasmic reticulum, for example, is upregulated to support elevated protein secretion by mature B cells through activation of the unfolded protein response (Shaffer 2004, Taubenheim 2012). Similarly, mitochondrial volume increases to support elevated respiration in myocytes following prolonged exercise (Jornayvaz and Shulman 2010). Peroxisome and lipid droplet volumes increase upon cellular exposure to environments enriched in very long chain fatty acids (Smith and Aitchison 2013, Thaim 2013). The importance of controlling organelle size is further suggested by the many scaling relationships that have shown both fixed relative sizes of various organelles compared to their host cells in a variety of organisms and developmental contexts (Levy and Heald 2012), including for the nucleus (Levy and Heald 2010, Chen 2019) and vacuole (Chan 2016), and nontrivial relationships such as maximal mitochondrial activity in intermediate sized cells (Miettinen and Bjorklund 2016). Dysfunctional organelle size control has been linked to a wide variety of metabolic, developmental, and neurodegenerative disorders (Hatch 2013, Shahmoradian 2019, Mourelatos 1990, Chang 2017). Uncoordinated regulation of organelle size can lead to severe phenotypic defects, such as impaired *Chlamydomonas reinhardtii* motility in uncoordinated flagellar length control (McVittie 1972), inappropriately sized secretory vesicles due to variability in Golgi size (Ferraro 2014), and impaired metabolism due to defects in mitochondrial (Toda 2016) and peroxisomal (Waterham 2007) fission among others. What remains underexplored, as also highlighted by complementary work simultaneous to our own documenting the coupling between organelle size and fluctuations (Bauer 2020), is development of a quantitative understanding of the precision with which organelle size variability is controlled, particularly for organelles that exist in multiple copies per cell.

Drawing on a combination of the theory of stochastic processes and quantitative fluorescence imaging, we directly examine two questions: how precisely the does the cell control the sizes of its organelles and what, if any, overarching quantitative principles collectively describe the patterns of observed organelle sizes despite the vastly different molecular mechanisms that implement size control?

## RESULTS

### Organelle size control appears to be constrained by a limiting pool of building blocks

In order to decipher the quantitative principles governing organelle size control, we reasoned that we could use a mathematical model of organelle biogenesis to interpret endogenous stochastic fluctuations in organelle size. Our first task in building a mathematical framework to quantify organelle size control was to be able to distinguish between the three general limits organelle growth is thought to fall into (Marshall 2016). In the first limit, termed constant growth, organelle growth occurs at a constant rate. In the second limit, termed negative feedback control, the cell constrains organelle growth rates to drive them towards a target size. In the third limit, termed the limiting pool, we assume that organelle sizes are constrained by a limited pool of building blocks from which they are assembled. In each limit, organelle size is affected by both size-specific processes, such as growth and disassembly, as well as number changing processes such as fission and fusion (Rafelski 2008, Lowe 2007). We therefore derived a stochastic model of organelle biogenesis that tracks the joint probability distribution of organelle numbers and sizes in single simulated cells. In this model organelles can be created de novo, decay, undergo fission and fusion, grow in size, and shrink (**Fig.1A**). Using the Gillespie algorithm (Gillespie 1977) we solve our model for the three general limits that organelle growth can take. The simulation is performed by tracking the number and sizes of organelles in a single cell until they reach steady state. We repeat these simulations for 1000 cells and extract organelle number and organelle sizes from each simulated cell (**Fig. S1**).

**Fig. 1:**
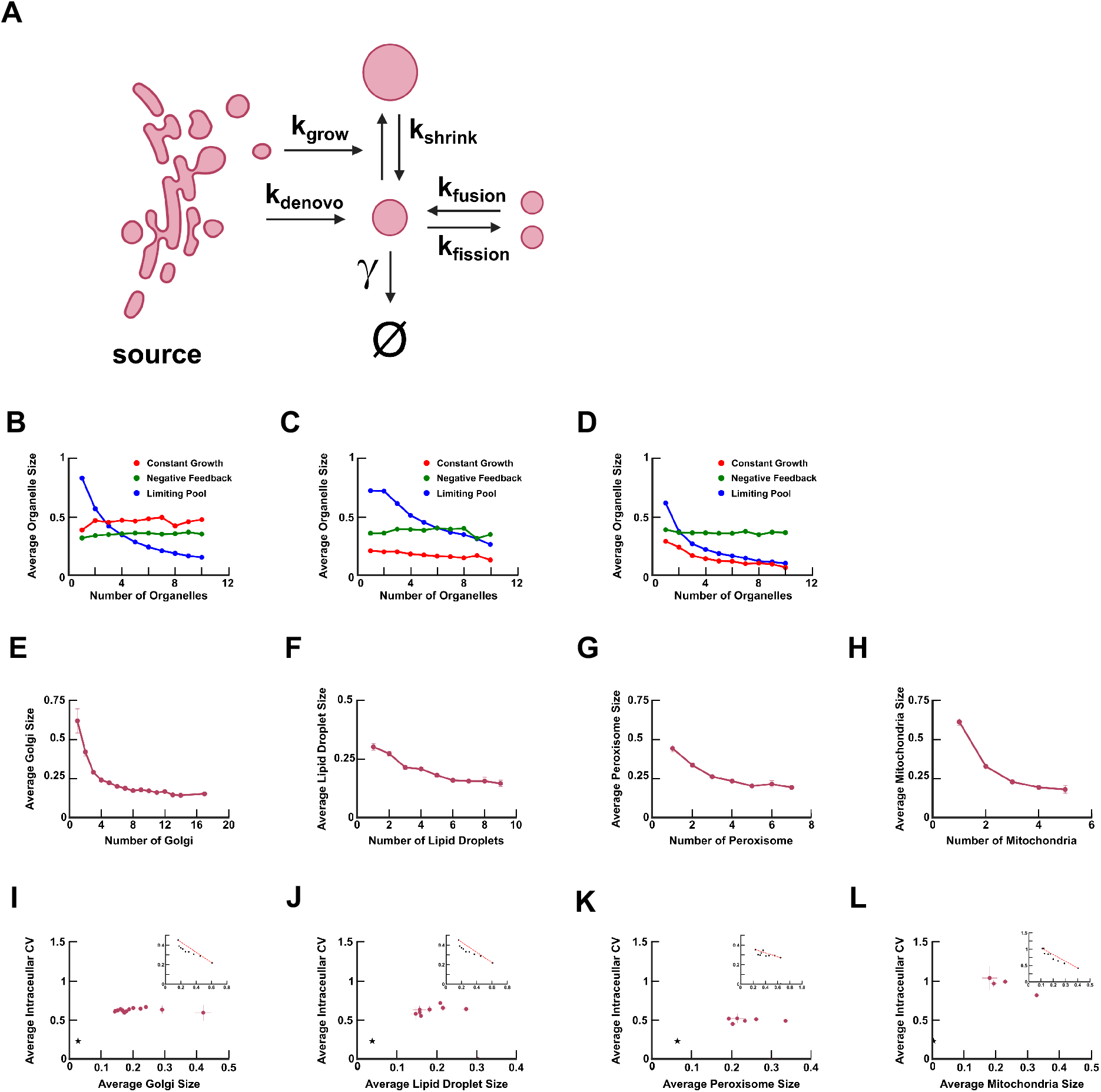
**A**) Schematic of mathematical framework. De novo synthesis of organelles, fission, fusion, organelle growth, shrink, and organelle decay occur at rate constants k_denovo_, k_fission_, k_fusion_, k_grow_, k_shrink_, and γ respectively. Mean organelle sizes vs. number of organelles from 1000 simulated cells at steady state in a regime where organelle abundances change through **B**) de novo synthesis and first order decay, **C**) de novo synthesis, fission, and first order decay, and **D**) fission and fusion. The red, green, and blue dots correspond to the constant growth, negative feedback, and limiting pool regime of our model. Experimental measurements of mean organelle size vs. number of organelles for fluorescently labelled **E**) Golgi body, **F**) lipid droplets, **G**) peroxisomes, and **H**) mitochondria. Average intracellular coefficient of variation (CV) of organelle sizes for **I**) late Golgi, **J**) lipid droplets, **K**) peroxisomes, and **L**) mitochondria; insets represent the predicted CV from our limiting pool model, and the blue stars correspond to the measured CV of diffraction-limited fluorescently labelled 100nm Tetraspeck microspheres.

Inspired by physiologically-relevant cases, we focused the simulations on three different regimes of number changing dynamics: in the late Golgi (Bevis 2002, Rossanese 1999) and lipid droplet (Pol 2014, Wilfling 2014, Walther 2017) relevant case in which number dynamics are governed by de novo synthesis and first order decay (**Fig. 1B**), in the peroxisome relevant limit of when organelle numbers change through de novo synthesis, first order decay, and fission (Hoepfner 2005, van der Zand 2012, Motley and Hettema 2007, **Fig. 1C**), and lastly in the mitochondria relevant limit of when the abundance of organelles changes solely through fission and fusion (Diaz 2008, **Fig. 1D**). For each number changing regime we see that the correlation between organelle number and average organelle size is diagnostic for whether organelles grow at a constant rate or are constrained by either negative feedback or a limiting pool of building blocks. Most importantly, in the limiting pool limit we observe a negative correlation between organelle number and average organelle size in both de novo synthesis and fission dominated organelle number controlling regimes; these correlations are robust to variations in the detailed parameters used in the simulation (**Fig. S2**). This contrasts with the constant growth regime, in which average organelle size is constant in simulated cells with different organelle numbers when organelles are made de novo. Intuitively, this is because in the constant growth regime with de novo synthesized organelles, the organelles within a given cell grow independently of each other and each attain a steady state size set only by the rates of assembly and disassembly irrespective of how many other organelles are in that given cell. Finally, we note that in the case of fission and fusion dominated organelle numbers, the constant growth and limited pool limits both reduce to the same picture as the fission and fusion processes conserve biomass, leading to an inherent tradeoff between organelle number and size. The model thus leaves us well positioned to use experimental data to infer which growth limit describes a given organelle.

To experimentally measure joint organelle number versus size distributions so as to infer organelle growth rules with our model, we analyzed the endogenous stochastic fluctuations in organelle numbers and sizes in the budding yeast *Saccharomyces cerevisiae*. To visualize the various organelles we examine, we fuse the fluorescent protein monomeric Kusibara Orange2 (mKO2) to organelle membrane resident proteins (**Fig. S3–5**). We obtain joint single cell average organelle size versus organelle number probability distributions of fluorescently labelled late Golgi (labelled with Sec7-mKO2), lipid droplets (Erg6-mKO2), peroxisomes (Pex3-mKO2), and mitochondria (Tom70-mKO2). From these joint distributions we plot the average organelle size as a function of organelle number. For each of the organelles we examine, we observe a significant negative correlation between organelle number and average organelle size (**Fig. 1E–1H**). In the case of late Golgi and lipid droplets, which we expect are dominated by de novo synthesis, the negative correlation is inconsistent with the constant growth and negative feedback limits of the general stochastic growth model. In the case of the mitochondria, whose copy numbers are a result of fission and fusion, the observed negative correlation is inconsistent with negative feedback. In the case of the peroxisome, the negative correlation is consistent with both constant growth with fission and the limiting pool limits. In order to distinguish between these pictures, we note that reducing peroxisome fission should flatten the curve relating organelle size to number if peroxisome growth occurs in the constant growth limit while it should remain negatively correlated in the limiting pool limit. We genetically deleted the peroxisome fission factors *DNM1, FIS1,* and *VPS1* (Kuravi 2006, Mukherji 2014, **Fig. S6A-C**) and observe the persistence of a negative correlation between average organelle size and organelle number. Our measurements suggest that the sizes of the four organelles under study are all constrained by a limited pool of building blocks.

### Cells exhibit robust average intracellular variability in organelle size

Among the most attractive hypotheses arguing for the utility of the limiting pool model of organelle growth is that it achieves a stable organelle size in the absence of feedback. However, it has been shown theoretically that growing multiple organelles from a limited pool of building blocks can lead to remarkably severe size fluctuations between organelles within the same cell (Mohapatra 2017, Fai 2019), potentially impairing cellular-scale physiological function. To quantify how intracellular fluctuations in organelle size behave in our model, we plot the average coefficient of variation (CV) in organelle size as a function of organelle size. We focus on cells simulated in the limiting pool limit of the model. As expected for a Poisson-type process, we observe that for decreasing organelle size (which corresponds to increasing organelle number) the intracellular organelle size CV increases. This suggests that cells face a fundamental tradeoff in their ability to achieve organelle size homeostasis. We then use our experimental joint distributions of organelle number and size for the late Golgi, lipid droplets, mitochondria and peroxisomes to directly measure the CV of organelle sizes within single cells (**Fig. 1I-L, Fig. S7**). Contrary to our theoretical expectation (**Fig. 1I-L**, insets), we see that the average intracellular CV of the late Golgi, lipid droplets and peroxisomes remain constant with varying average organelle size. Thus, despite organelles changing in size by 2-fold, cells are able to robustly maintain the fluctuations in sizes to within a factor of 0.5 (peroxisomes) to 0.6 (late Golgi and lipid droplets) of the mean organelle size, thereby avoiding a rise in intracellular fluctuations when the mean organelle size shrinks. Only the average intracellular CV of mitochondria appears to increase when the cell creates more, smaller copies of this organelle.

### Burst-like organelle growth model derived to explain robust organelle size control unifies description of endomembrane organelle size statistics

To address the discrepancy between the noise profile resulting from our model versus the noise profile of organelles observed experimentally, we consider a fundamental revision to how organelle growth proceeds in our mathematical framework. Building on previous observations that subcellular structures can grow from bursts of random sizes of building blocks (Ludington 2013) we constructed a model in which organelles grow from a limited pool of building blocks in exponentially distributed bursts of random size β that occur at random times characterized by a burst frequency α (**Fig. 2A**). In the model, changes in organelle size arise from modulation of the burst size β, which is proportional to the free pool of building blocks available for organelle growth.

**Fig. 2:**
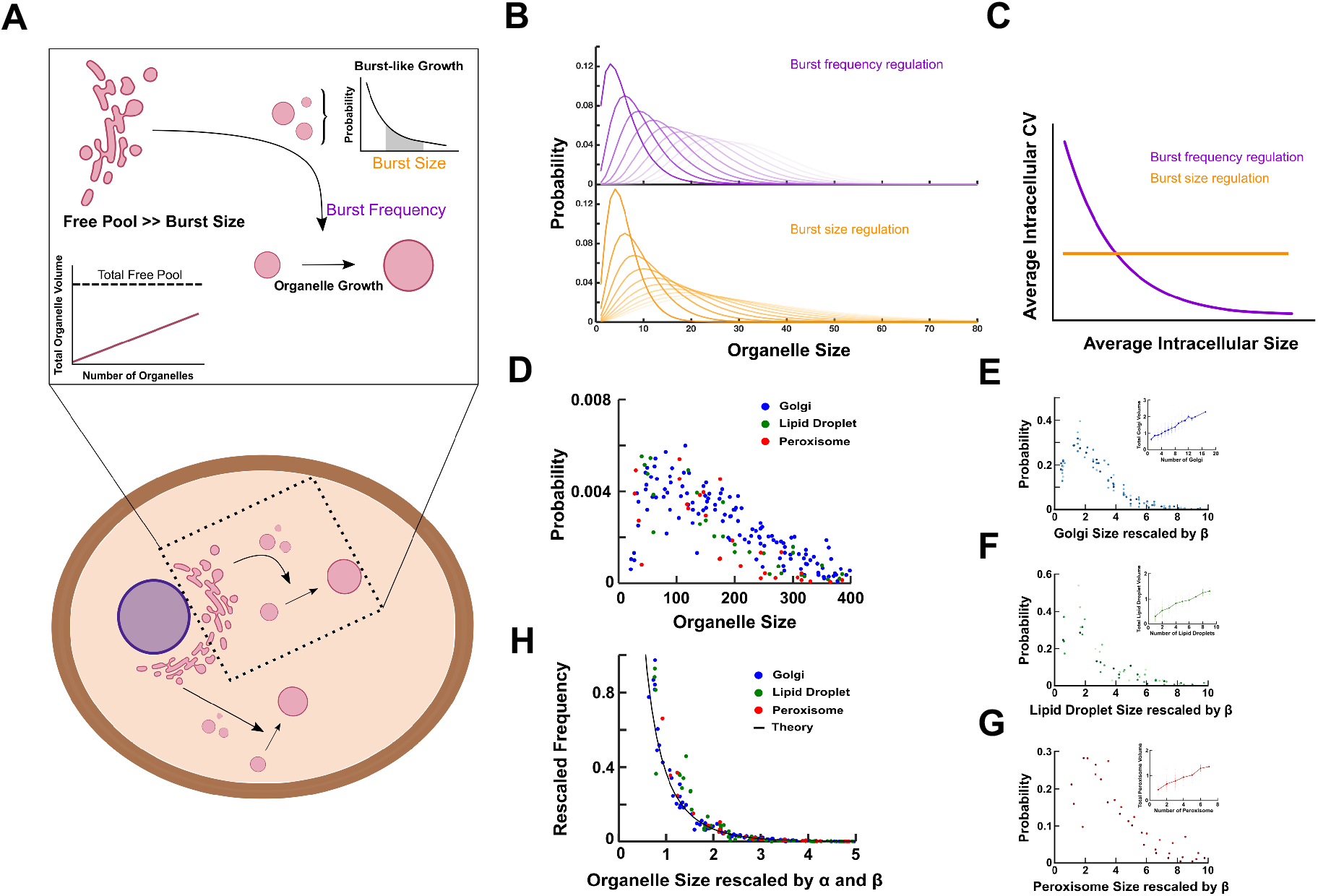
**A)** Schematic depicting model of burst-like growth in organelle size in the limit of large number available of building blocks for organelle growth compared to the typical size of growth bursts. **B)** Organelle size distributions predicted from burst-like organelle growth model under either modulation of burst frequency (purple histograms) or burst size (orange histograms). **C)** Organelle size fluctuations as a function of mean organelle size under burst frequency (purple curve) or burst size (orange curve) modulation. **D)** Histograms of experimentally measured late Golgi, lipid droplet, and peroxisome sizes. **E)** Histograms of late Golgi, **F)** lipid droplet, and **G)** peroxisome sizes from (**D**) rescaled by their experimentally derived organelle-specific burst sizes β; insets are the corresponding total organelle size as a function of number of organelles. **H)** Histogram of organelle size distributions rescaled by both experimentally derived organelle-specific burst sizes β and burst frequencies α; black curve is the theoretically predicted scaling relationship from the model, *f(s) = e^−s^/s*.

If the limited pool of building blocks is not exhausted and allows organelle sizes to fluctuate independently of each other, and if copy number changes are slow compared to size changes, then the resulting organelle sizes from cells that contain a defined number of organelles follow a gamma distribution with mean size αβ (**Fig. 2B**). Crucially, the average intracellular organelle size fluctuations depend only on the burst frequency α but is independent of the burst size β (Friedman 2006). This allows the cell to decouple the average fluctuations in intracellular organelle size from the mean organelle size (**Fig. 2C**) as we observe experimentally, allowing robust tuning of organelle sizes.

The model predicts that if the pool of building blocks is not depleted, we should observe two types of data collapses upon rescaling organelle sizes (**Fig. 2D, Fig. S8A-G**). First, since the model holds that organelle sizes change only through modulation of burst sizes, rescaling organelle sizes by their corresponding burst sizes should collapse them onto unifying gamma distributions specific to each organelle. Second, by further rescaling these organelle-specific size distributions by their organelle-specific burst frequencies, our whole collection of late Golgi, lipid droplet, and peroxisome sizes should collapse onto the universal curve *f(s) = e^−s^/s*, where *s* is the dimensionless rescaled organelle size (**Methods**).

To test our predictions from the burst-like organelle growth model we first construct a measure of pool depletion. To measure pool depletion, we measure the total volume of all copies of a given organelle in each cell, and assume all cells have roughly equal amounts of the limiting pool. Plotting the total organelle volume as a function of organelle number, if the total volume increases linearly with the number of organelles, we conclude that the pool has not depleted. However, if at higher organelle numbers the total organelle volume plateaus then we conclude that the limited pool has depleted. We plot the total organelle volume versus organelle number for late Golgi, lipid droplet, and peroxisomes (**Fig. 2E-G, insets**). We observe that while each organelle appears to grow from a limited pool, the pool of building blocks does not appear to deplete.

To test our prediction that modulation of burst size alone can explain changes in organelle sizes, we first use our experimental data to obtain burst size values for each late Golgi, peroxisome and lipid droplet size distribution. Mathematically, the burst size is the product of the mean organelle size and the square of the average intracellular CV, both of which we measure experimentally. We then rescale late Golgi, peroxisome, and lipid droplet sizes by their experimentally inferred burst sizes and plot the resulting rescaled size distribution. We see that late Golgi, lipid droplet, and peroxisome sizes collapse onto single gamma distributions (**Fig. 2E-G**). Further rescaling the organelle-specific rescaled size histograms by their experimentally calculated burst frequencies α, we see that despite the starkly different molecular mechanisms by which their sizes are controlled, late Golgi, lipid droplet, and peroxisome size distributions further collapse onto the single theoretically predicted universal curve *f(s) = e^−s^/s* (**Fig. 2H**). This fitting parameter-free data collapse strongly suggests that our mathematical model of organelle biogenesis has captured an essential, unifying feature of endomembrane organelle growth regulation.

### Depletion of limited pool of building blocks breaks robust organelle size control

According to our model, sufficient capacity in the limited pool of building blocks allows organelles to grow and fluctuate in size independently of each other and allow for robust intracellular size control. Our model also predicts, however, that if the available amount of building blocks for growth depletes enough to be similar to the typical burst size (**Fig. 3A,B**) then intracellular organelle size fluctuations will become anti-correlated. Anti-correlated fluctuations, in turn, will lead to a rise in the average intracellular CV (**Fig. 3C**).

**Fig. 3:**
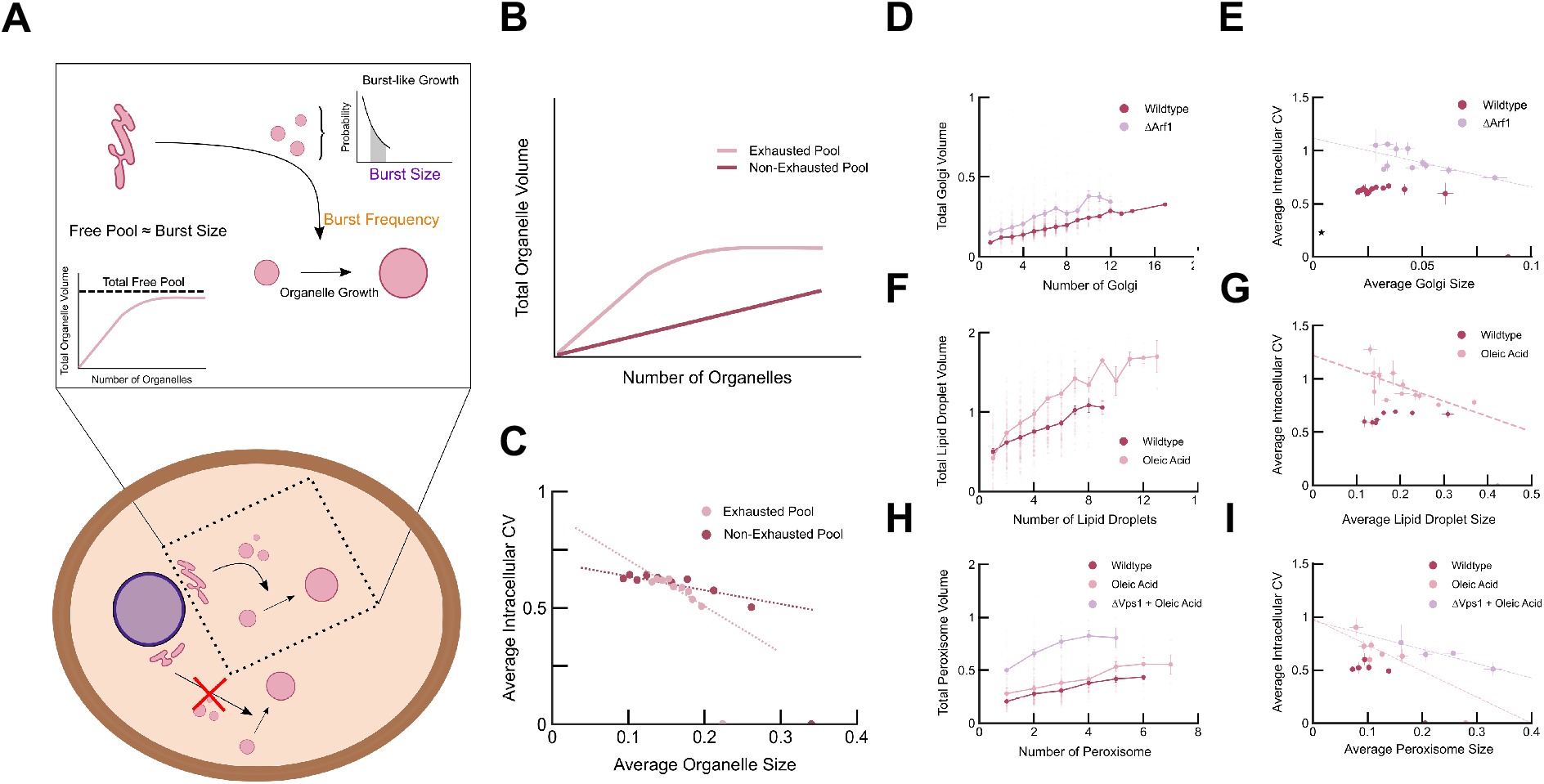
**A)** Schematic depicting model of burst-like growth in organelle size in the limit of a similar number of available building blocks for organelle growth compared to the typical size of growth bursts. **B)** Theoretical prediction of total organelle size V as a function of the number of organelles in the limits where the pool is exhausted (pink line) versus non-exhausted (maroon line) **C)** Average intracellular CV of organelle sizes in 1000 simulated cells from burst-like model, when pool is not depleted versus when pool is depleted. Total organelle size **D)** and **E)** average intracellular size CV of late Golgi from populations of wildtype (maroon) and *ΔARF1* (purple) cells. **F)** Total organelle size and **G)** average intracellular size CV of lipid droplets from populations of wildtype cells grown in glucose (maroon) and in medium with 0.2% oleic acid (pink). **H)** Total organelle size and **I)** average intracellular size CV of peroxisomes from populations of wildtype cells grown in glucose (maroon) and of wildtype (pink) and *ΔVPS1* (purple) cells grown in medium with 0.2% oleic acid.

To test our prediction that organelle growth from a limited pool with no spare capacity leads to an elevated intracellular CV, we examine the case of mitochondria, whose sizes are a balance between biomass conserving fission and fusion. We plot total organelle volume vs. organelle number for mitochondria and observe a flat line (**Fig. S9A**), consistent with the idea that cells largely rearrange an existing a fixed pool of mitochondrial biomass when changing mitochondrial number through fission and fusion. The resulting organelle size distributions are consistent with our model (**Fig. S9C-I**) and with our observation of an increasing intracellular CV with decreasing mitochondria size (**Fig. 1L**).

Next we test our prediction that downregulating size growth bursts will increase average intracellular size fluctuations. We reason that inhibiting Golgi vesicular traffic could reduce large bursts of size changes, leading to a narrower distribution of burst sizes that would mimic deterministic growth steps, or reduce the burst frequency. Both changes would lead to depletion of the available supply of building blocks and a predicted increased average intracellular Golgi CV. To test this idea, we analyze the late Golgi in cells lacking the vesicular traffic regulator *ARF1* (Bhave 2014). We observe, in agreement with our hypothesis, that upon deletion of *ARF1* the average intracellular Golgi size CV increases as the average Golgi size decreases (**Fig. 3D,E**). Moreover, we observe a concomitant increase in cell size variability upon increased Golgi size variability (**Fig. S10**).

Finally, to test our prediction that pool depletion leads to an elevated intracellular organelle size CV, we turn to lipid droplets and peroxisomes. Lipid droplets and peroxisomes are dynamic organelles whose sizes and copy numbers increase upon cellular exposure to long chain fatty acid-rich environments (Wilfling 2014, Walther 2017, Mast 2010). We hypothesized that culturing cells in an oleic acid rich environment could expose capacity constraints in lipid droplet and peroxisome biogenesis given our previous observation that these organelles appear to grow from a limited pool of building blocks. We observe that when grown in glucose, neither lipid droplets nor peroxisomes deplete the limited pool of building blocks, however, when grown in medium rich in oleic acid, total organelle volume plateaus (**Fig. 3F,H**), suggesting that the pool has depleted. Concomitantly, we observe that the lipid droplet and peroxisome average intracellular organelle size CV increases with decreasing mean organelle size (**Fig. 3G,I**). Importantly, we observe the elevation of the average intracellular peroxisome size CV in oleic acid even upon deletion of the primary peroxisome fission factor *VPS1*, (**Fig. 3I**) suggesting that capacity constraints from the limited pool of building blocks, and not elevated fission, are responsible for the increased noise in peroxisome sizes.

### Pattern of organelle size robustness is conserved to human induced pluripotent stem cells

To establish the evolutionary conservation of our observed pattern of robustness in organelle size, we leveraged single cell imaging data from human induced pluripotent stem cells (iPSC) whose Golgi apparatus and mitochondria are labelled available from the Allen Cell Atlas (Roberts 2017). With single cell organelle size distributions from populations of iPSCs, we are able to construct datasets of both average organelle size versus organelle number and the average intracellular organelle size CV as we have done for budding yeast. Both the Golgi apparatus and mitochondria yield negative sloping average organelle size versus organelle number curves consistent with a limited pool model constraining their growth (**Fig. 4A,C**). Most significantly, we observe that the average intracellular organelle size CV exhibits the same pattern in human iPSCs as seen for budding yeast: a robust, invariant organelle size CV as a function of average organelle size for the Golgi (**Fig. 4B**) and a sensitive, inverse correlation between organelle size CV and average organelle size for the mitochondria (**Fig. 4D**).

**Fig. 4:**
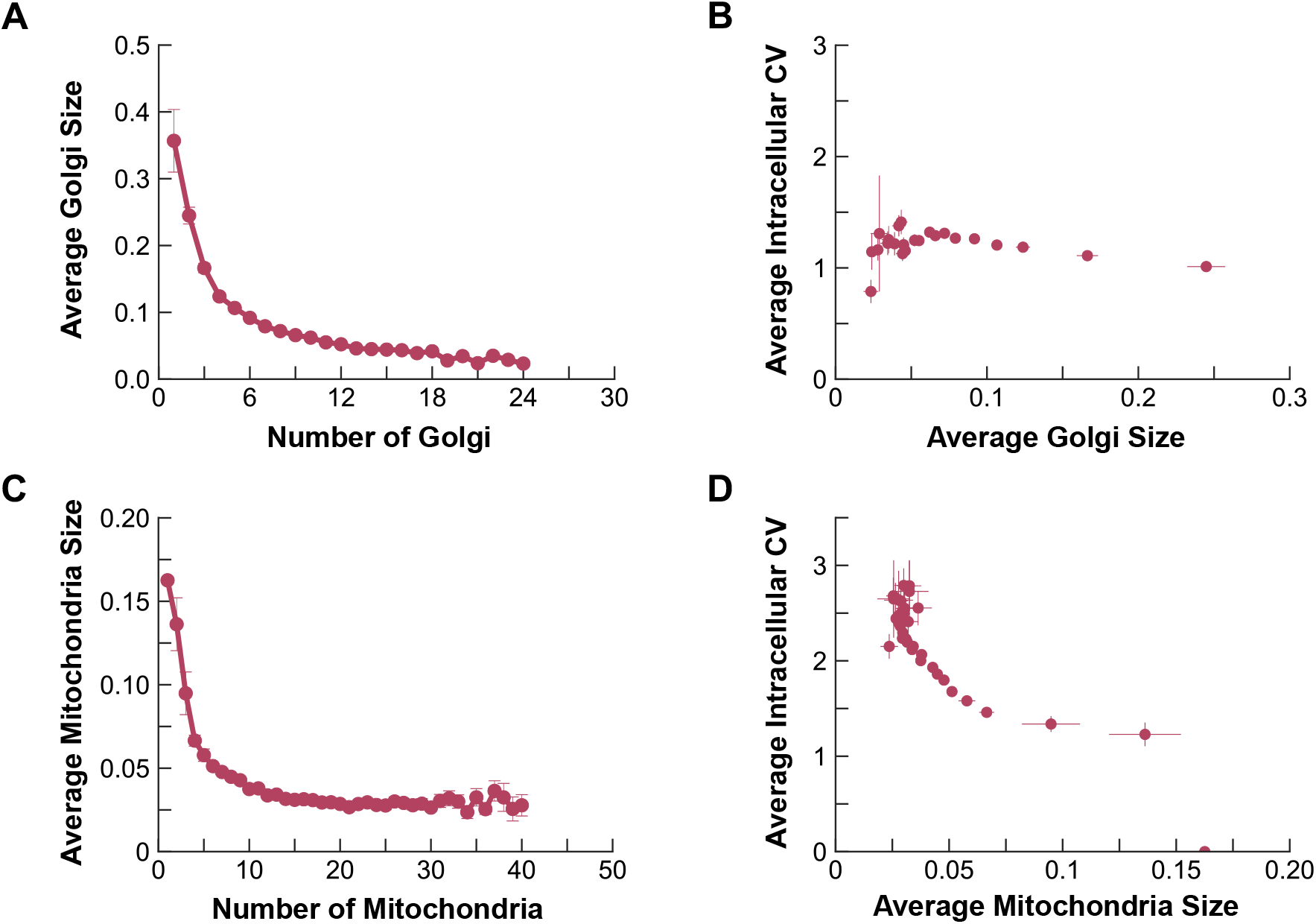
**A)** Average Golgi size versus number of Golgi from images of single induced pluripotent stem cells taken from the Allen Cell Atlas. **B)** Average Golgi intracellular size CV versus average Golgi size. **C)** Average mitochondrial size versus number of mitochondria from images of single induced pluripotent stem cells taken from the Allen Cell Atlas. **D)** Average mitochondrial intracellular size CV versus average mitochondrial size.

## DISCUSSION

In order to explain our observed invariance of average organelle size fluctuations to changing mean organelle size, we propose a model in which organelle growth proceeds in a burst-like fashion from a limited pool of building blocks. The pattern of organelle size robustness is shared between budding yeast and human iPS cells. The underlying molecular mechanisms producing these bursts are yet to be fully elucidated and are likely to be organelle-specific and potentially species-specific: Golgi size, for example, is likely influenced by both small increases in size from non-vesicular traffic as well as the sudden, large increases in size from vesicle fusion as we have shown here, while lipid droplet size bursts may result from burst-like expression of genes such as those encoding neutral lipid synthesis enzymes. However, the size statistics of a diverse array of organelles appear to be well described by a single unifying model that can be used to interpret future studies on the mechanistic underpinnings of organelle size control and a basis for more sophisticated modeling efforts to more accurately capture details suppressed here (Craven 2016, Choubey 2019, Banerjee and Banerjee 2020) as well as comparison to universal growth phenomena in seemingly unrelated biological systems (Iyer-Biswas 2014). More generally, the robust organelle size control we observe here may play a significant role in cellular homeostasis and aid in efforts to engineer organelle size to rationally control cellular metabolism. Suppression of average intracellular organelle size variability in principle allows the organelles within the cell to be functionally interchangeable with each other, avoiding fluctuations in phenotype that could arise from variability from one organelle to another within the same cell arising solely due to increased number of that organelle. Establishing the limits of the ability of the cell to suppress size fluctuations, which we demonstrated in the case of Golgi, peroxisomes and lipid droplets and complementary work in flagellar length control in *Chlamydomonas reinhardtii* showing a coupling between flagellar length and length fluctuations (Bauer 2020), may provide important clues for how to design organelles to control cellular phenotypes. Our results point to the potentially general principle that that robust size control can, remarkably, be the result of random burst-like assembly of subcellular structures.

## ACKNOWLEDGEMENTS

We thank D. Laman Trip, T. Maire, and H. Youk for critically reviewing our manuscript. K.A. was supported by a fellowship from the Center for Science and Engineering of Living Systems at Washington University in St. Louis.

## AUTHOR CONTRIBUTIONS

S.M. conceptualized the project. K.A. performed the computational simulations and imaging experiments, assisted by A.K. A.K. analyzed the image data, assisted by K.A., and prepared the manuscript figures. K.A., A.K., and S.M. wrote the manuscript.

## COMPETING INTERESTS

None declared.

## METHODS

### Mathematical framework for growth with deterministic step sizes

In our mathematical framework (**Fig. 1A**), solve for the dynamics of the probability of a state of organelles in a single cell that undergo the following changes:

I. De novo synthesis of organelles from a given source of constituents within the cell with a constant rate k_denovo_

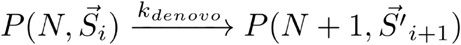
II. Decay of organelles due to inheritance, maturation, autophagy, etc. at a rate γ per organelle per time:

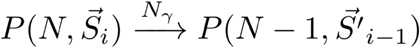
III. Creation of two organelles from a single organelle through fission at a rate k_fission_ per organelle per time:

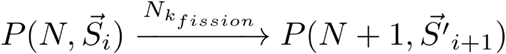
IV. Creation of a single organelle through the fusion of two organelles at a rate k_fusion_ per organelle per squared time:

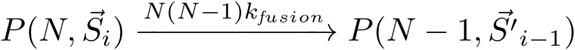
V. Organelle size growth at a rate r_growth_ per time:

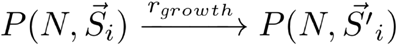
VI. Organelle size decay at a rate k_decay_ per number of organelle constituents (size s) per time:

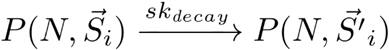

We initially assume that organelles grow in deterministic increments (packet sizes). In all three limits, we assume that the rates of decay, fission, fusion, and shrinking are Nγ, Nk_fission_, N(N−1)k_fusion_, and S_i_k_shrink_ respectively, in which N is the number of organelles and S_i_ is the size of the i^th^ organelle and S^’^ refers to the size of each organelle in the new probability state. In the case of constant growth, we assume that the growth and de novo synthesis rates are equal to their rate constants respectively (i.e., r_denovo_ = k_denovo_ and r_grow_ = k_grow_). In the negative feedback model, we assume that the de novo synthesis rate is equal to its rate constants, however the growth rate is constrained by a feedback term

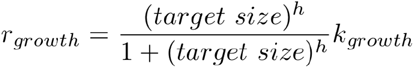

in which target 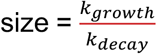 and h is the Hill coefficient. Lastly, in the kdecay limiting pool model we characterize the rate for de novo synthesis and organelle growth as R_denovo_ = k_denovo_N_pool_ and R_grow_ = k_grow_N_pool_ respectively, in which N_pool_ is the free pool available for the organelles. Additionally, we assume that the pool is replenished by the process of organelle decay and shrinking.

In **Fig. S1A-F** we plot the trajectories of a single cell reaching steady-state for the three limits of our framework, in a regime where the abundance of organelles change through de novo synthesis, fission, and first order decay, and in a regime in which organelle abundance changes solely through fission and fusion.

To obtain additional insight about the three limits of our model, we sweep through a large parameter space and simulate 1000 cells. From each simulation we obtain the slope of the plot of average organelle sizes vs. organelle abundance. Finally, we plot the histogram of the recorded slopes by increasing fission rate (purple represents a fission rate of zero and red represents the highest fission rate in our simulations). Comparing the slope histograms, we observe a zero slope for almost all parameter sets in the negative feedback model and a negative slope for all parameter sets used to simulate the limiting pool model. However, we notice that in the constant growth model, the slopes become increasingly negative as fission rates increase (**Fig. S2G–2I**). We implement this information to learn about peroxisomal growth. In wildtype yeast, we consider peroxisomes to undergo de novo synthesis, fission, and first order decay. Our measurement of the wildtype data results in a negative correlation between average size and abundance of peroxisomes (**Fig. 1**). However, this can only help us rule out the negative feedback control model, as we expect a negative correlation between number and average organelle size at high fission rates in the constant growth model. In order to determine the correct model to describe peroxisomes, we engineered strains of yeast with their peroxisomal fission factors deleted, while grown in medium rich in long chains of fatty acid. Peroxisomes undergo fission when grown in oleic acid. We observe that even in the strain with the *VPS1* gene deleted (*VPS1* being the quantitatively most significant of all fission factor deletes, Ref. 1), the correlation between the abundance and average peroxisome size is still negative in **Fig. S6**. This argues against constant growth as the underlying mechanism for peroxisomal growth, suggesting that peroxisomes grow from a limited pool of building blocks.

### Experimental methods

#### Strains

*Saccharomyces cerevisiae* strain BY4742 was obtained as a kind gift from H. Zaher. Cells were transformed by standard lithium acetate methods to fluorescently labelled indicated organelles with monomeric Kusibara Orange 2 (mKO2) obtained from Addgene (Cambridge, MA). Peroxisomes in strains engineered with deletions of *DNM1, FIS1,* and *VPS1* were visualized as in Ref. 1.

#### Culture conditions

For glucose medium, strains were grown to mid-log phase at 30°C in standard synthetic medium containing 2% glucose and subsequently imaged. For oleic acid medium, strains were grown to mid-log phase at 30°C in standard synthetic medium containing 2% glucose, washed twice, and resuspended in medium containing 0.3% yeast extract, 0.6% peptone, 0.1% glucose, 0.1% Tween40, and 0.2% oleic acid, cultured for 20 hours in the oleic acid rich medium, and subsequently imaged.

#### Imaging Conditions

Wildtype strains were imaged with a Nikon Ti2 microscope using a Hamamatsu Orca Flash v3 scientific CMOS camera with standard TRITC emission and excitation filter cubes. The exposure time for the samples ranged from 50 - 250ms. Strains engineered with deletions of *DNM1, FIS1,* and *VPS1* were imaged as in Ref. 1.

#### Image Processing

Custom Matlab code was written to segment, filter, binarize, and compute sizes of all organelles. Cellular segmentation was semi-automated and varied between datasets. For the confocal images, segmentation was performed by hand with ImageJ. For the camera images, cell segmentation was performed by labeling the cellular cytoplasm with YFP, whose image was binarized to identify those pixels belonging to each cell. Following cell segmentation, 3D images of organelles were extracted from the field of view to be measured individually and deconvolved (**Fig. S3**). Two thresholding processes were used, dependent upon the morphology of the organelle. For the globular structures (lipid droplets, peroxisome, and Golgi) the filtering began with a weak Gaussian blur followed by taking the Laplacian of the image. Ultimately, the threshold for binarization was picked by hand and confirmed by eye. For the tubular structures (mitochondria) we used the Hessian based “Frangi Vesselness” filter (available on mathworks.com); the threshold was again picked by hand and confirmed by eye. After applying the filters corresponding to the given structure’s morphology (globular or tubular), the organelle images from each cell are binarized one z-slice at a time (**Fig. 1B**). Z-slices are then reconstructed to form a 3D image of the cell and built-in Matlab functions are used to identify individual organelles and count their voxels (**Fig. S3**).

To test the robustness of the analysis pipeline to choice of threshold, we perform two analyses. First we show that the measured average intracellular CVs for a population of the given organelle type where Threshold 2 is 10% greater than Threshold 1 are highly correlated between the thresholds (**Fig. S7**). Second, we observe no correlation between the average pixel intensity of a given organelle and its size, eliminating a possible source of artefactual size variation due to differences in fluorescent protein concentration; shown here is data on the late Golgi (**Fig. S4**). Finally, to test that our spatial resolution following deconvolution was sufficient to detect variation in organelle size and not be truncated by diffraction, we measured the fraction of organelles in our dataset that were indistinguishable in size from a diffraction limited, 100nm Tetraspeck bead (Thermo Fisher T7279), whose image data was processed in the same way outlined in Supplementary Fig. 6, with the modification of being carried out without the cell segementation step. For the images collected with a camera, we report the proportion of organelles from all cells that fall at the diffraction limit (**Fig. S5**). In the vast majority of cases, this proportion is below 10% (**Fig. S5**, numbers above individual violin plots).

### Mathematical framework for growth with burst-like step sizes

In the revised mathematical framework describing burst-like steps of organelle growth, the number changing processes (Equations I-IV above) and the organelle shrink process (Equation VI above) from the initial model remain unchanged. The growth process has two modifications: first the rate of growth is given as a constant “burst frequency” parametrized by α, second the growth increment is itself a random quantity described by an exponential distribution parametrized by a “burst size” β. As shown by Friedman et al. (Ref. 2), the steady state distribution of organelle sizes resulting from these stochastic processes, assuming that the number dynamics are slow compared to the size dynamics, is a gamma distribution:

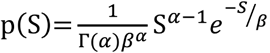

In the model, we note the burst size itself varies according to how much free pool of building blocks is available for growth; namely β = β_0_N_pool_. Intuitively, therefore, an increased number of organelles will lead to a smaller average organelle size as the fraction of building blocks held in organelles and away from the pool is greater, thereby reducing the burst size.

In order to collapse the data onto organelle-specific gamma distributions, we can rescale the organelle sizes by their corresponding burst sizes. We define a new random variable

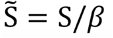

Then by the chain rule:

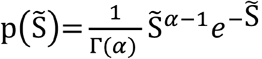

where each organelle-specific gamma distribution is parametrized by its organelle-specific burst frequency alpha. In order to further collapse all organelle data onto a universal master curve, we can rescale 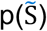 by multiplying by a factor of 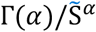 to remove any dependence of the right-hand size on α. The resulting scaling relationship is the dimensionless curve plotted in **Fig. 2H**:

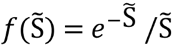

#### Figure schematics

Figure schematics in Fig. 2A and Fig. 3A were performed using images from BioRender.

## FIGURE SUPPLEMENTS

**Fig. S1.**
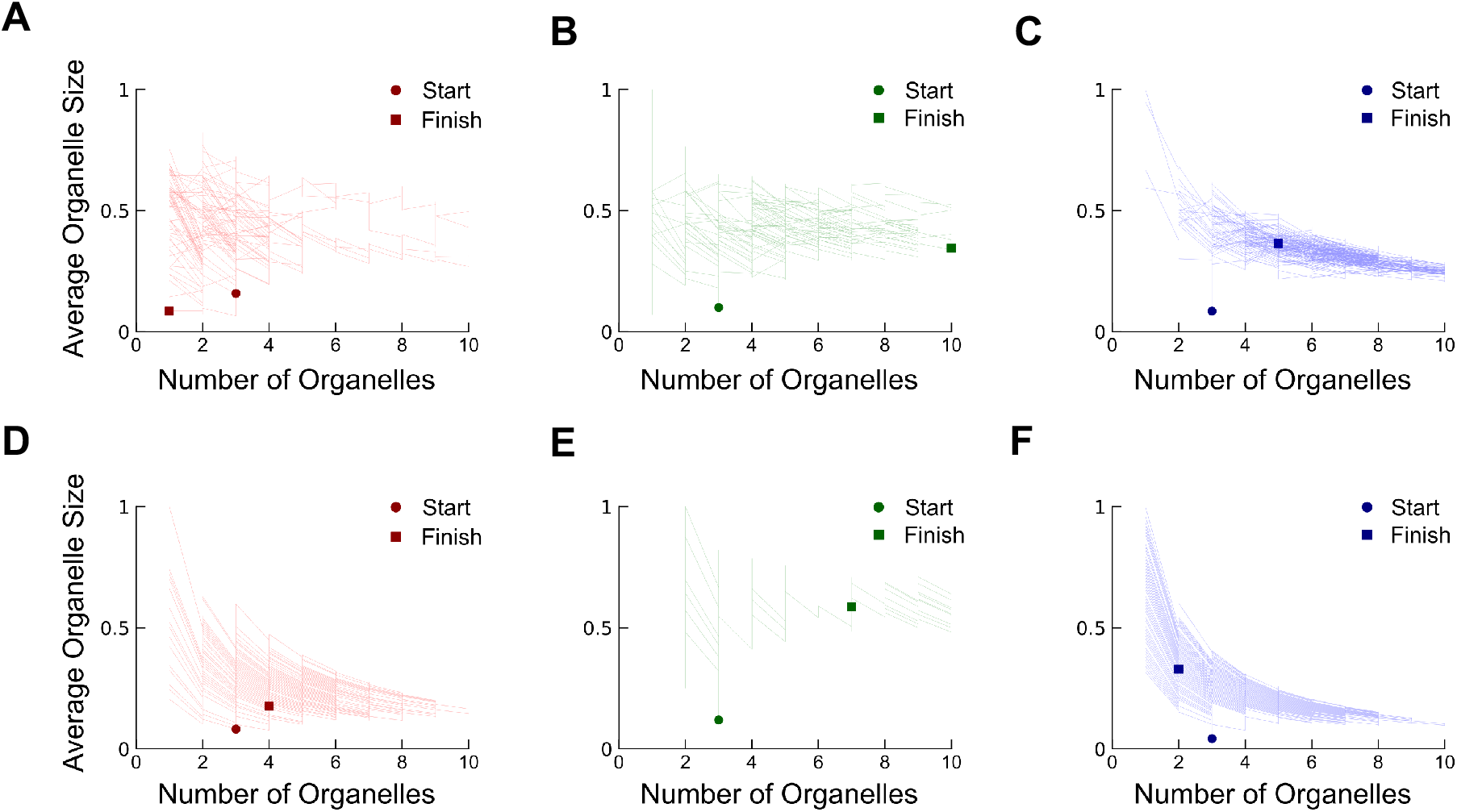
In the regime where organelle abundances change through de novo synthesis, fission and first order decay **A**) trace of a simulated single cell in the limit of constant growth **B**) negative feedback control, and **C**) limiting pool, from t=0 (circle) to steady-state (square). In the regime where organelle abundances change through fission and fusion **D**) trace of a simulated single cell in the limit of constant growth **E**) negative feedback control, and **F**) limiting pool, from t=0 (circle) to steady-state (square).

**Fig. S2.**
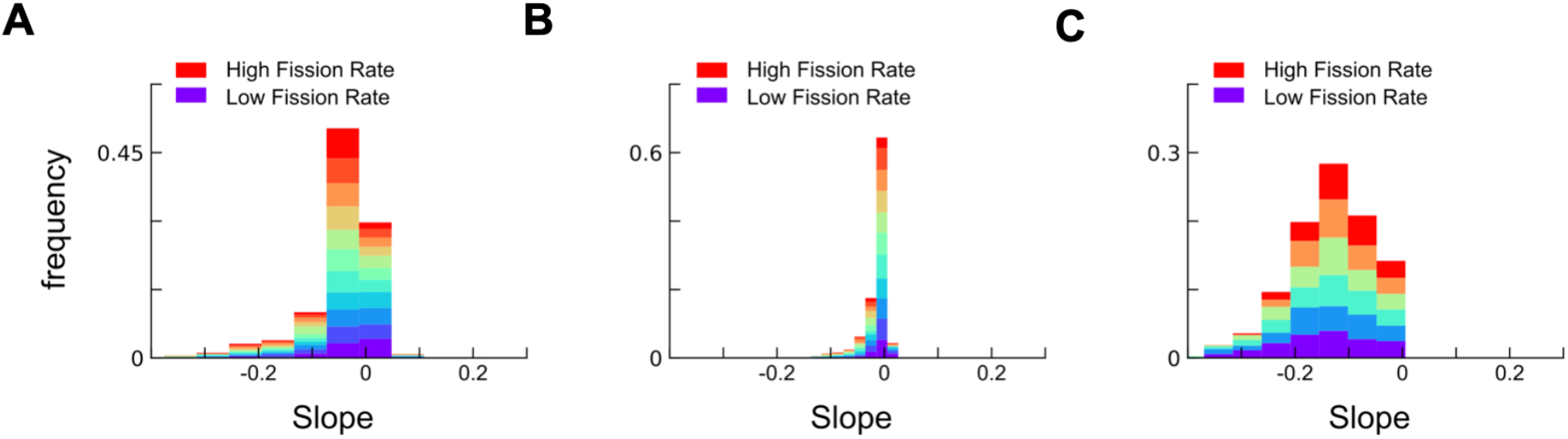
Histograms of slopes relating average simulated organelle size and organelle number obtained from sweeping through a large parameter space from kfission = 0 to high fission rates for **A**) the constant growth, **B**) negative feedback, and **C**) limiting pool model.

**Fig. S3.**
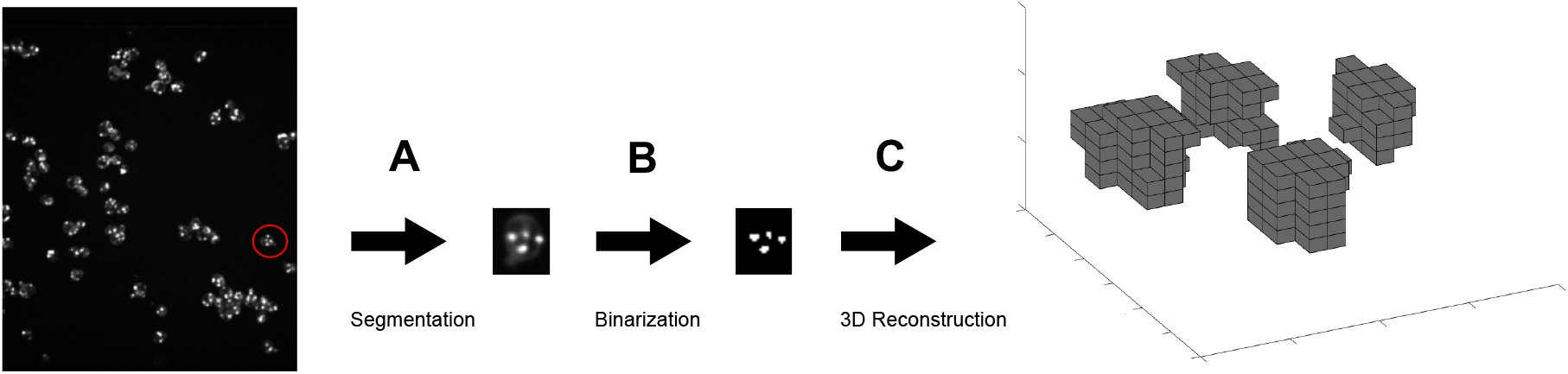
**A**) After semi-automated segmentation, individual cells are extracted from a given field of view. **B**) After applying the filters corresponding to the given structure’s morphology (globular or tubular), each cell is binarized one z-slice at a time. **C**) Z-slices are then reconstructed to form a 3D image of the cell and built-in Matlab functions are used to identify individual organelles and count their voxels.

**Fig. S4.**
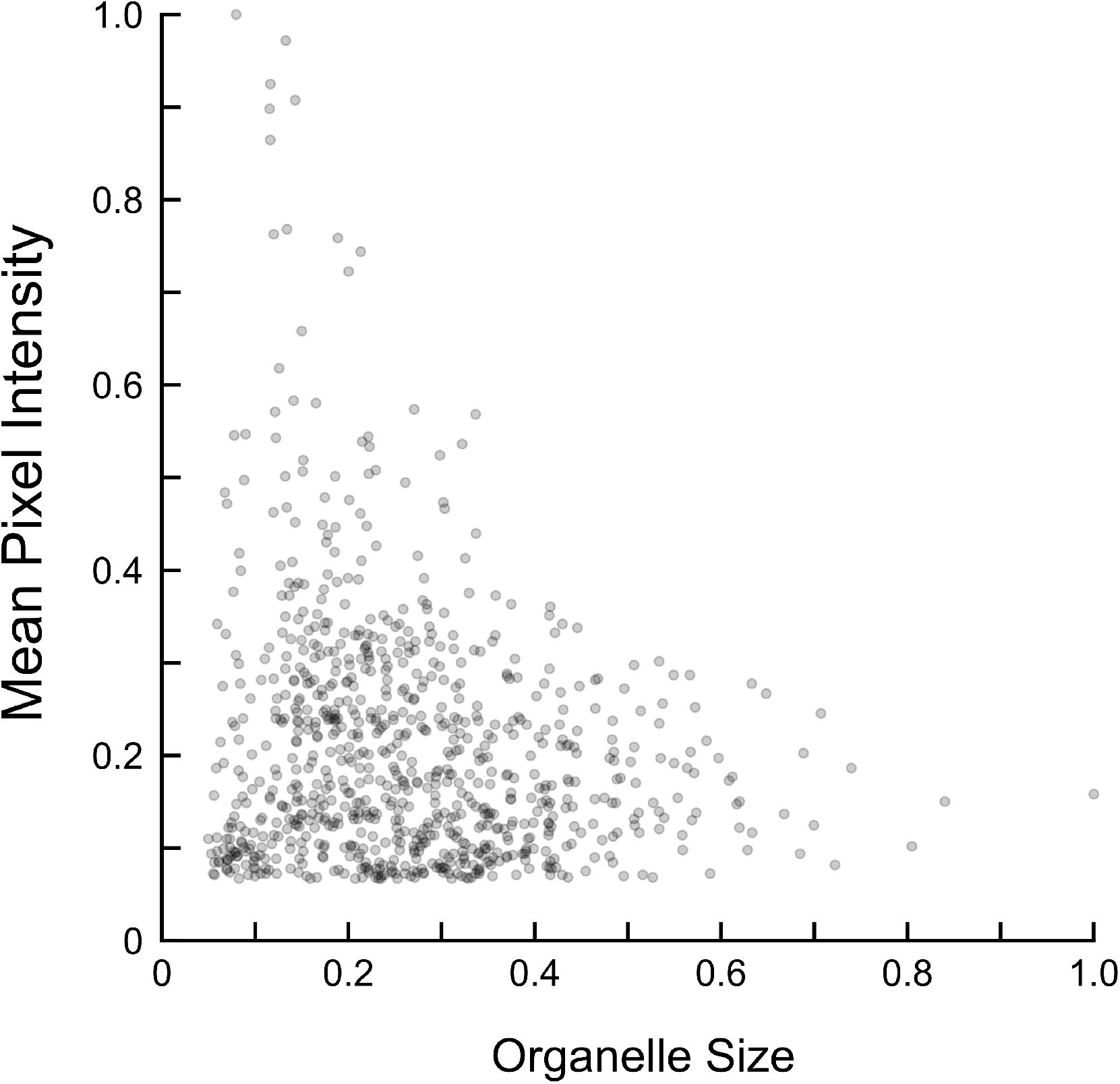
Scatter plot of late Golgi, labelled by Sec7-mKO2, average pixel intensity versus measured late Golgi size.

**Fig. S5.**
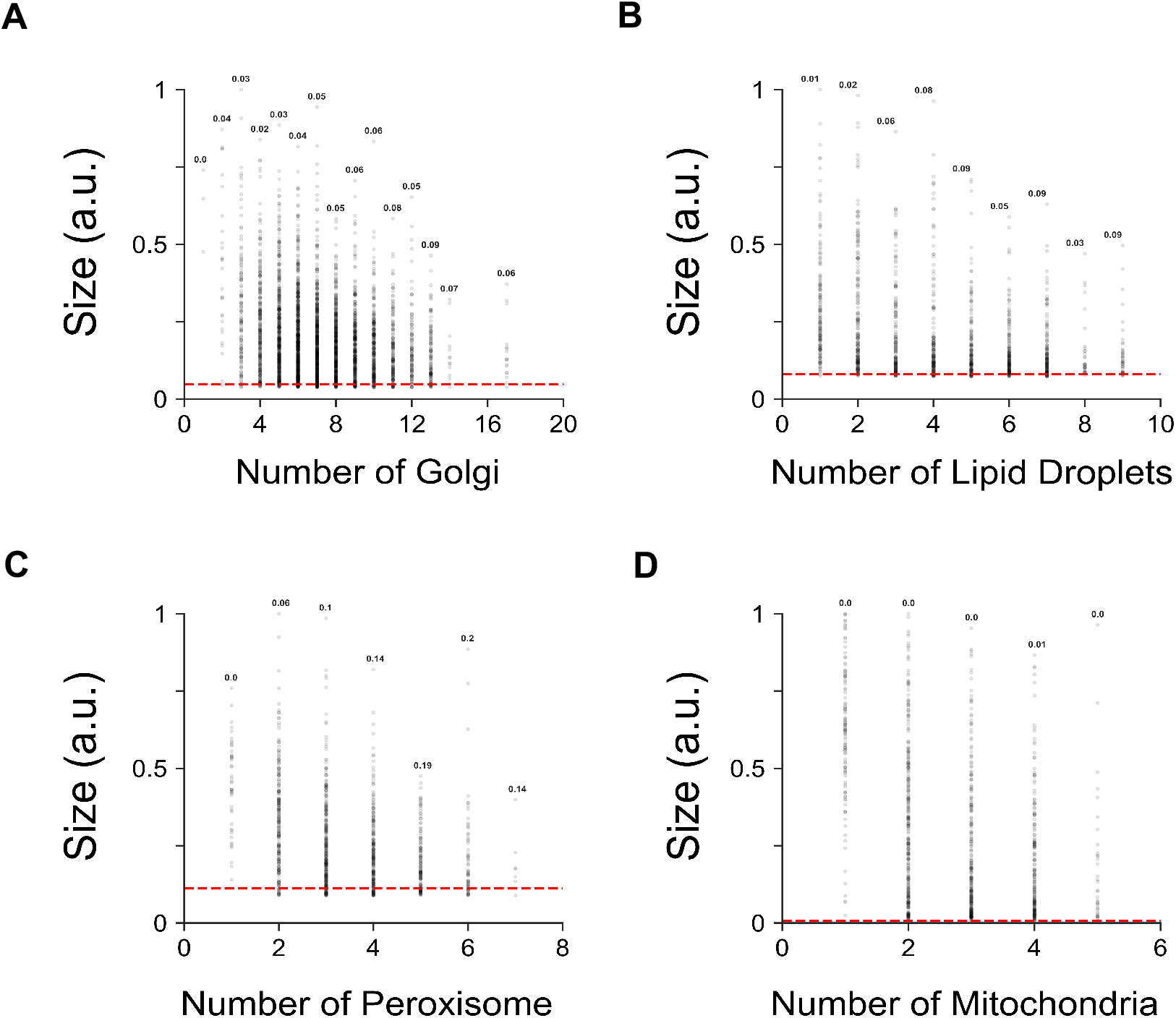
Proportion of organelles fall at the diffraction limit (computed by measuring the size of a diffraction-limited 100nm Tetraspeck microsphere bead). Floating numbers above data points are proportion of organelles from cells in a subpopulation with defined number of organelles that falls at the diffraction limit.

**Fig. S6.**
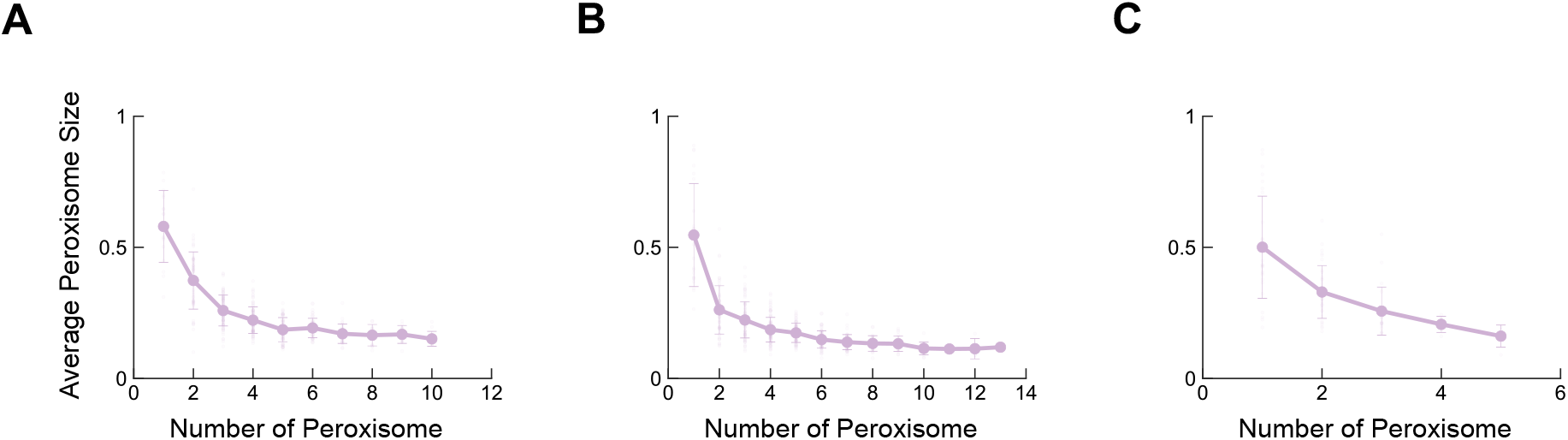
Average of mean peroxisome sizes as a function of peroxisome number in strains grown in medium rich in oleic acid with their peroxisomal fission factors **A**) *DNM1*, **B**) *FIS1*, and **C**) *VPS1* deleted.

**Fig. S7.**
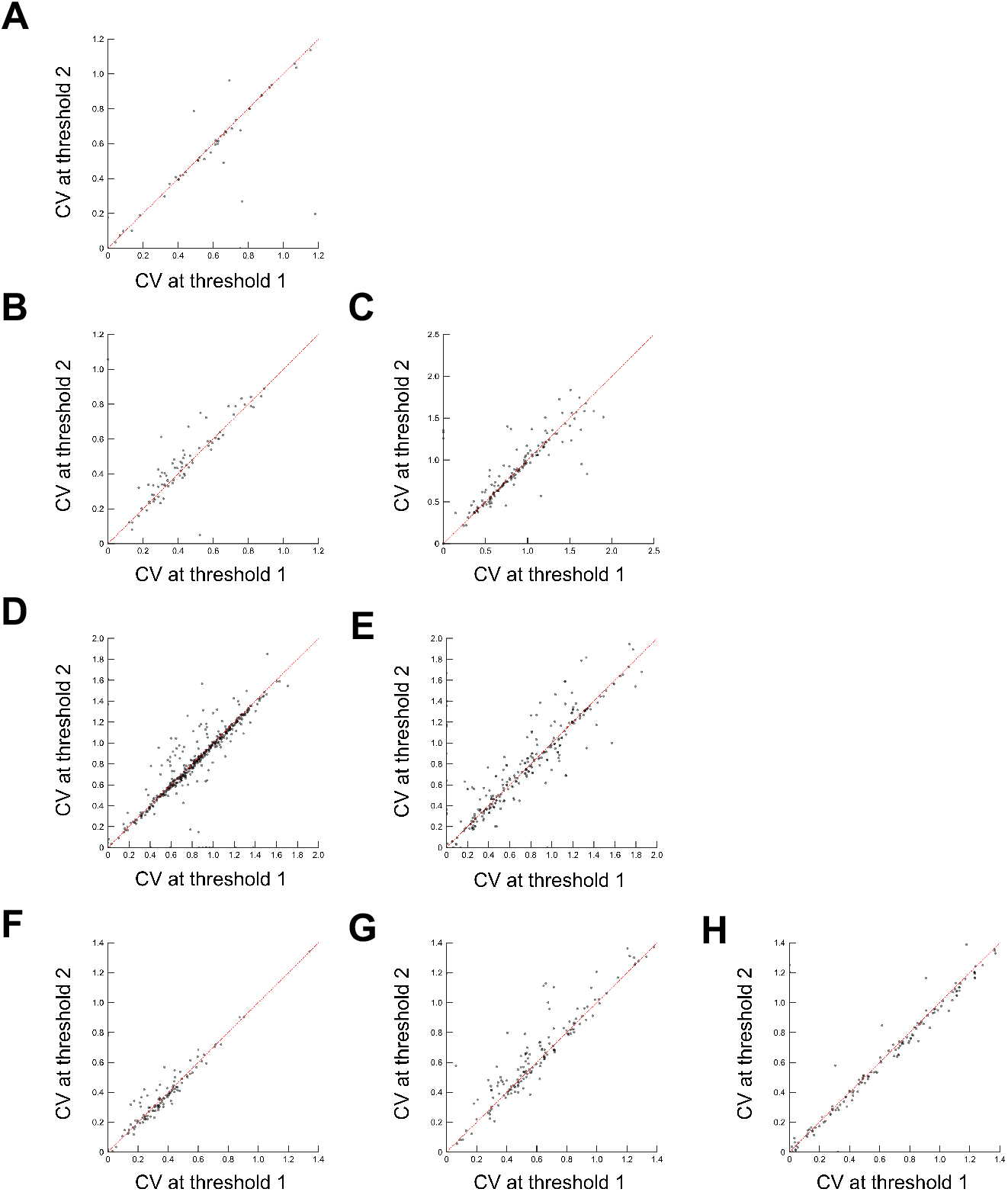
Sensitivity analysis of measured average intracellular coefficient of variation to choice of threshold. Threshold 1 is the threshold used for image analysis whose results are presented in Fig. 1, 2, and 3. Threshold 2 is a 10% higher pixel intensity threshold. **A**) Tom70-mKO2, **B**) Sec7-mKO2, **C**) Sec7-mKO2 Δ*ARF1*, **D**) Erg6-mKO2, **E**) Erg6-mKO2 (oleic acid), **F**) Pex3-mKO2, **G**) Pex3-mKO2 (oleic acid), **H**) Pex3-mKO2 Δ*VPS1*

**Fig. S8.**
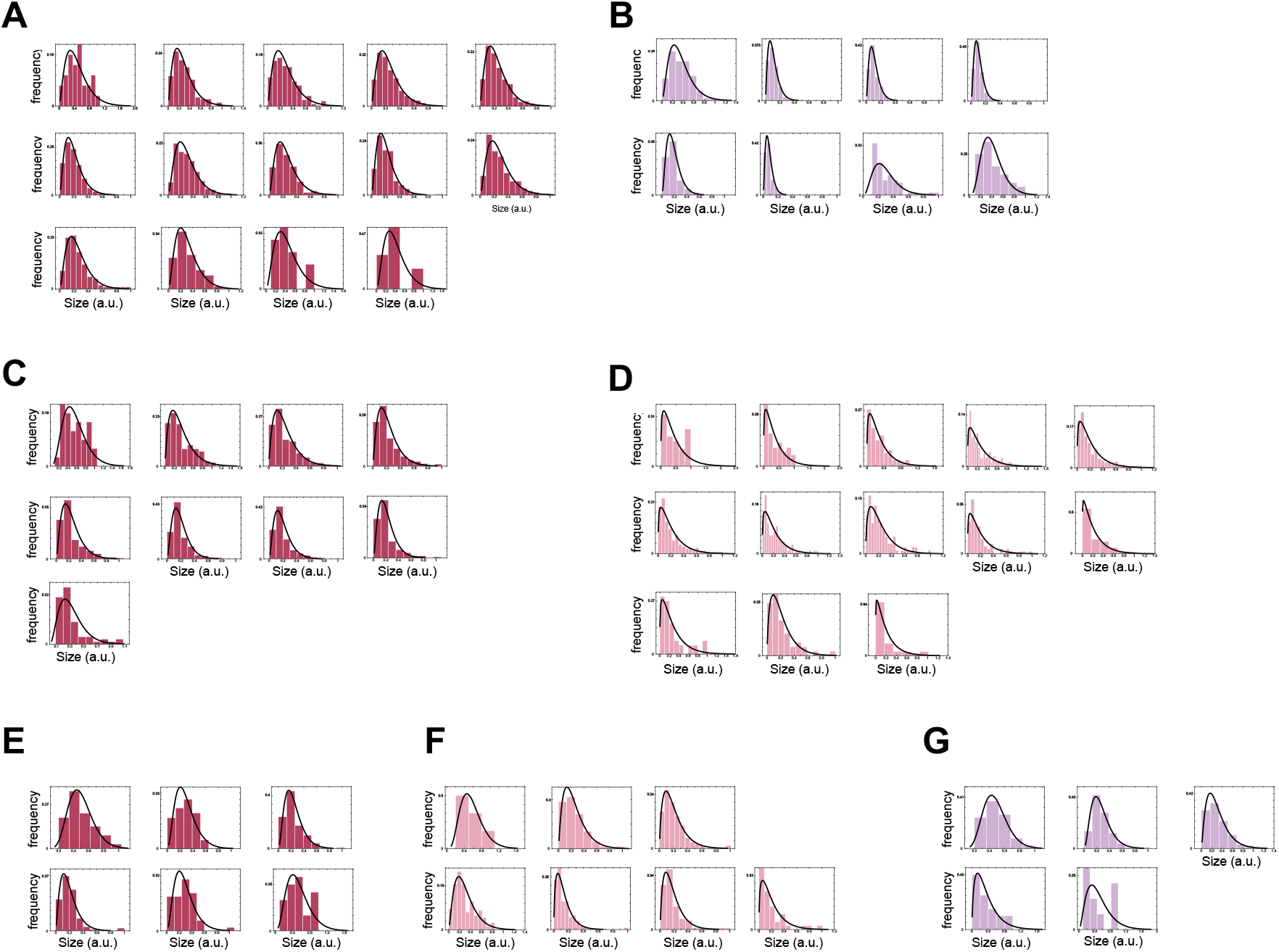
Single organelle size distributions and corresponding gamma distribution fits to the **A**) late Golgi apparatus in Sec7-mKO2 strains, **B**) late Golgi in Sec7-mKO2 strain with *ARF1* deleted, **C**) lipid droplets in Erg6-mKO2 strains grown in glucose, and **D**) in medium rich in oleic acid, **E**) peroxisomes in Pex3-mKO2 strains grown in glucose, **F**) in medium rich in oleic acid, and **G**) strain with *VPS1* deleted and grown in medium rich in oleic acid.

**Fig. S9.**
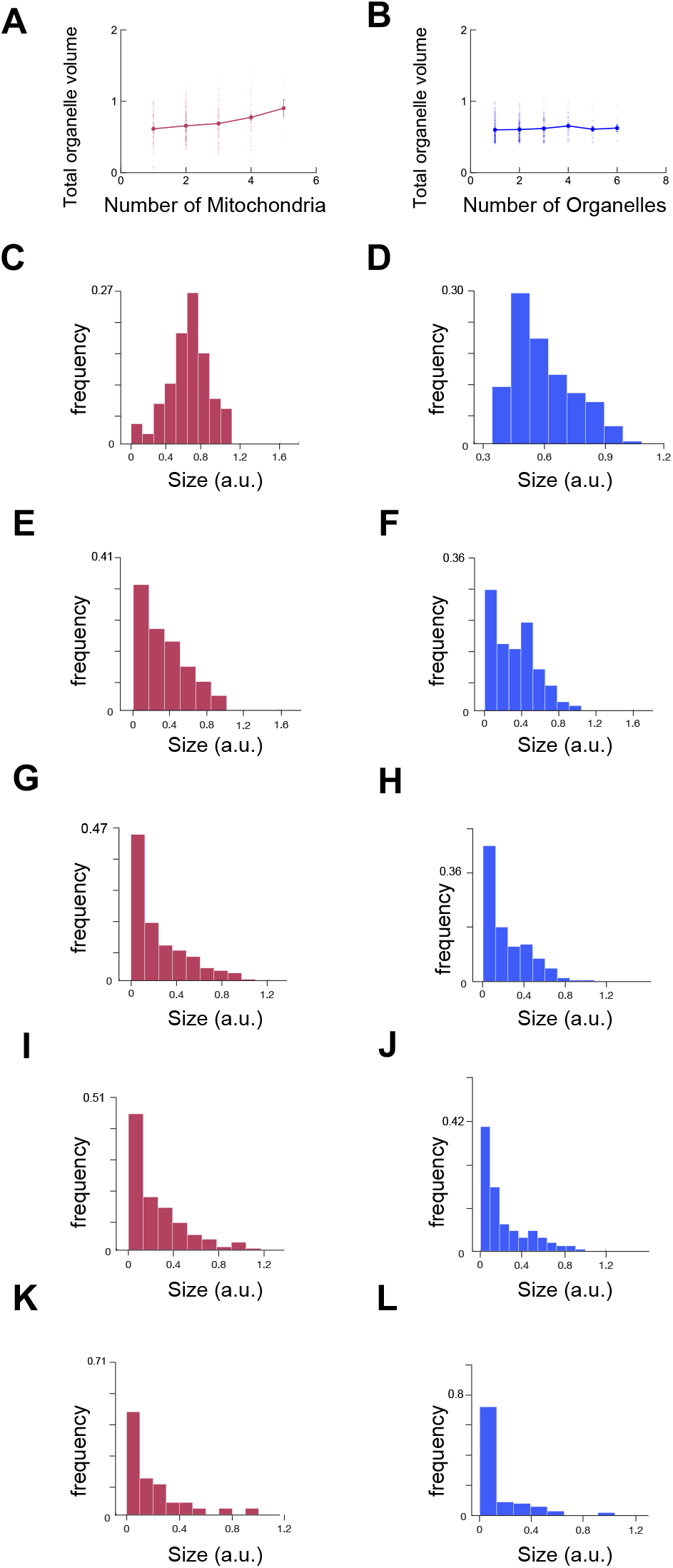
Total organelle volume vs. number of organelles plot of **A**) mitochondria (red) and **B**) result of bursty model (blue) with organelle numbers changing through fission and fusion. **C**-**L**) distribution of mitochondria (red) and simulated organelles (blue) sizes from subpopulations of cells with defined mitochondrial copy numbers.

**Fig. S10.**
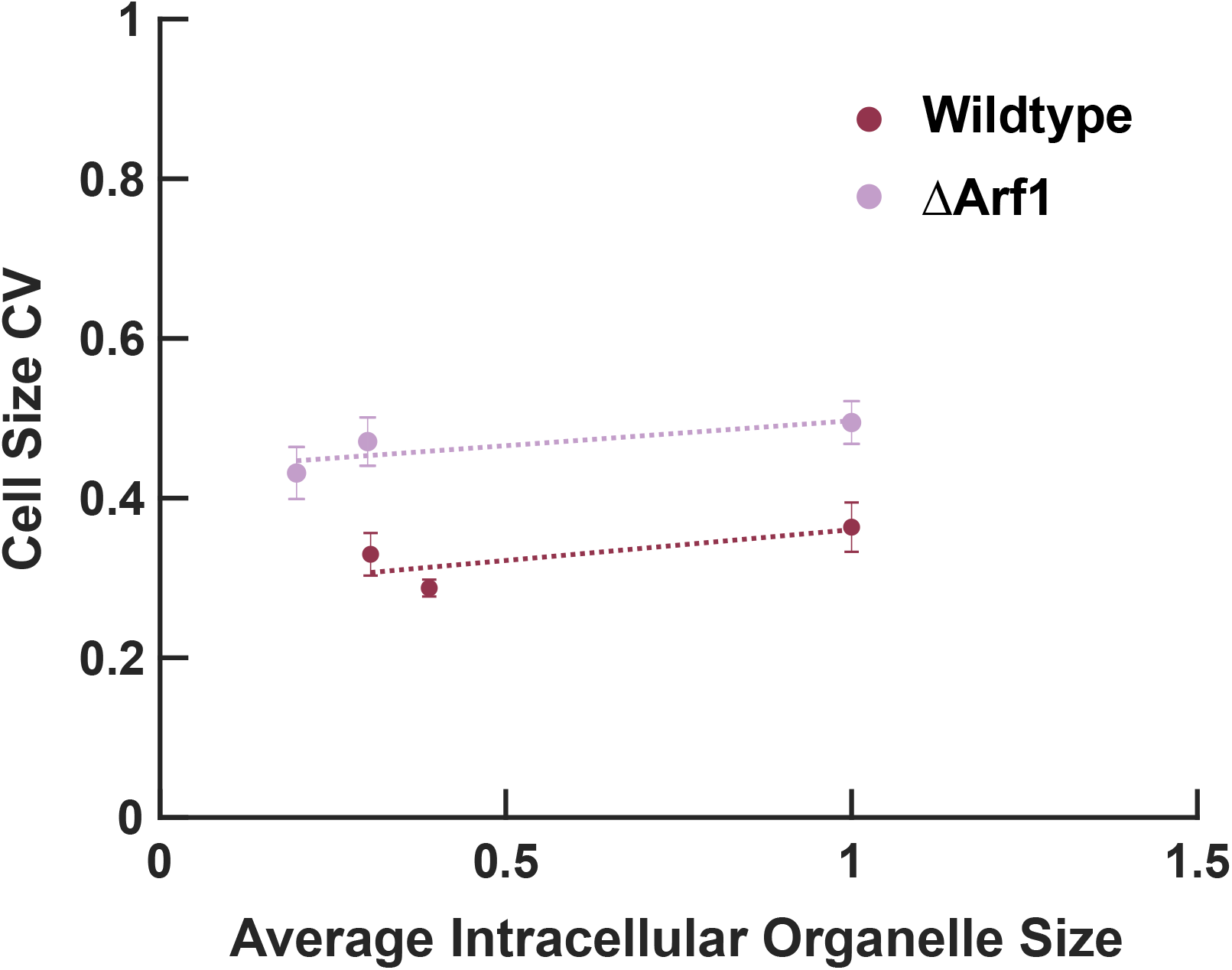
Comparison of cell size coefficient of variation in the wildtype strain versus a strain carrying a deletion of *ARF1*, which exhibits an increased average intracellular coefficient of variation in late Golgi sizes. Cells were binned into 3 quantiles according to their average late Golgi size, their sizes were estimated using their area as measured by a YFP fluorescent marker for their cytoplasm, and the CV was computed from the resulting cell size distributions.

## Notes

### Competing Interest Statement

The authors have declared no competing interest.

### Summary of Updates

Data-driven rescaling method to test model with no fitting parameters used. Extension of findings to Golgi and mitochondria of mammalian iPS cells.

